# Alteration of neural network and hippocampal slice activation through exosomes derived from 5XFAD nasal lavage fluid

**DOI:** 10.1101/2023.01.24.525465

**Authors:** Sang Seong Kim, Jae Kyong Jeon, Dulguun Ganbat, Taewoon Kim, Kyusoon Shin, Sungho Hong, Jong Wook Hong

**Author notes:** Correspondence to: Jong Wook Hong, Hanyang University, Gyeonggi-do & Seoul, Republic of Korea, Sang Seong Kim, Hanyang University, Gyeonggi-do, Republic of Korea.

## Abstract

Exosomes contain various intracellular biomarkers reflecting the condition of cells, organs, and subjects. Under neurodegenerative conditions, they contrive in detrimental neuronal communications leading to the initiation and propagation of neurodegenerative symptoms. Since the exosomes in olfactory fluid are readily accessible to collect in ample amount noninvasively and highly enriched in neural biomarkers, they can be a primary biomarker if we can verify precise pathophysiological information from them. Here we show that exosomes from nasal lavage fluid (NLF) of the neurodegeneration model animal, 5XFAD mice, induce the pathological network activity in neuronal cultures. We separated intact NLF exosomes from the 5XFAD and wild-type mice via a high-efficacy microfluidic system and applied them to the primary cortical neurons and organotypic hippocampal slice cultures (OHSC), whose neuronal activities were monitored by a high-density microelectrode array system. We found that NLF exosomes from 5XFAD mice increased the firing rate of neuronal spikes with augmentation of neuronal connectivity similar to the effect of pathological amyloid beta oligomer treatment. Furthermore, the current source densities, computed from the local field potentials, were elevated in OHSCs incubated with the exosomes, suggesting a pathological shift in synaptic and membrane currents. Those results demonstrate that NLF exosomes from neurodegeneration model can effectively modify neuronal networks and suggest that this property can serve as a functional biomarker for Alzheimer’s disease.

## Introduction

In Alzheimer’s disease (AD), neuronal death in the hippocampus and the entorhinal cortex leads to cognitive impairment and memory loss. Although the pathogenic mechanisms of AD have not been completely elucidated, increases in amyloid oligomer and hyperphosphorylated Tau in the brain of AD patients have been reported (Braak and Braak 1991; Devanand et al. 2007; Serrano-Pozo et al. 2011; Thal et al. 2002). Several factors, such as mitochondrial dysfunction, oxidative stress, endothelial dysfunction, and autophagy dysfunction, have been found to contribute to synaptic loss between neurons, which can lead to neural network destruction (Moreira et al. 2009; Reddy and Beal 2008). With the progression of AD, the severity of dementia symptoms may vary according to the modality of pathogenesis. During the mild cognitive impairment (MCI) stage, there is an evident impairment in memory, reasoning, and sensory information processing (Teixeira et al. 2012). As a result, functional imaging of the brains of MCI patients frequently indicates epileptic instability (Allen et al. 2007; Dickerson et al. 2005). Furthermore, neuronal recordings in AD model animals showed alternating tonic and phasic response patterns followed by an elevation in calcium concentration (Busche et al. 2008; Busche and Konnerth 2015; Verret et al. 2012). It is unclear to what extent the underlying cause of neuronal hyperactivity in AD can be attributed (Busche et al. 2012; Grienberger et al. 2012; Wang et al. 2006). Currently, it is thought that surrounding Aβ42 stimulates neurons through receptor irritation or that Aβ42 uptake leads to excitation through unknown mechanisms (Zhong et al. 2018; Zotova et al. 2010). Regardless, a complete understanding of such neuronal hyperactivity prior to cellular breakdown in AD is still elusive.

Typically, the AD symptoms manifest once it has reached a fatal stage with no chance of reversible recovery of neurons. Patients often miss opportunities for recovery due to the period of time between the onset of disease and the development of symptoms. Hence, early detection of AD is essential to decelerating its progression and providing time for therapeutic intervention. There is no doubt that cerebrospinal fluid (CSF) examination and PET imaging of the brain constitute the most convincing diagnostic calibers (Cohen et al. 2019; Klyucherev et al. 2022). For example, the presence of hyperphosphorylated Tau peptide (P-Tau), total Tau protein (T-Tau), and neurogranin in CSF provides over 90% accuracy in the diagnosis of AD (Palmqvist et al. 2017). Though both methods are effective, the invasiveness and clinical cost are critical caveats in practical settings. To avoid these defects, other methods are recommended to detect AD biomarkers in blood and other body fluids (Nakamura et al. 2018; Palmqvist et al. 2019). The credibility of fluid detection was previously subject to substantial variability due to the technical limitation of being able to detect very small amounts of biomarkers in comparison to CSF in less than 100 fold. Despite this limitation, though, it’s now possible to identify Aβ42 in blood down to subpicograms (Teunissen et al. 2022). As such, other body fluid biomarkers are being investigated in sophisticated ways using tear, urine, sweat, saliva, and nasal lavage fluid (NLF) with relative ease compared to the invasive methods (Jeromin and Bowser 2017; Klyucherev et al. 2022; Obrocki et al. 2020; Teunissen et al. 2022; Zetterberg and Bendlin 2021). It has been shown that NLF in particular contains amyloid precursor protein (APP) and α-synuclein reflecting the pathological changes that occur in AD, making it an efficient source of AD markers (Jung et al. 2022; Kovacs et al. 1996; Ohm and Braak 1987; Shi et al. 2014; Yoo et al. 2020). In the myriad of substances within NLF, exosomes are primarily categorized on the basis of their neuromodulatory effects on cellular interaction, synaptic plasticity, and signal transmissions (Ching and Kingham 2015; Frohlich et al. 2014; Saeedi et al. 2019; Sharma et al. 2019; Thery et al. 2002). Since nasal-derived exosomes can be used as diagnostic tools in various diseases symptoms, we sought to isolate exosomes from NLF in AD model animals and evaluate their functional effects on neurons. Among various types of AD models, 5XFAD was chosen because it overexpresses APP and presenilin 1 (PS1), which reliably induce pathogenic Aβ42 oligomer formation (Eimer and Vassar 2013; Forner et al. 2021; Oblak et al. 2021).

A majority of electrophysiological studies in AD model animals have been conducted in individual neurons to determine whether neuronal excitation or synaptic plasticity are impaired (Hu et al. 2018; Qiao et al. 2014). While such a method provides a sophisticated explanation of cellular physiology, it does not provide an overview of global changes in neuronal networks. The recording of multiple neurons in close proximity has been attempted using the paired clamping or multielectrode arrays, however, it was difficult to measure the activity pattern of neural networks. With the advent of high-density multielectrode arrays (HD MEA), thousands of neurons can be recorded simultaneously down to the subcellular level, allowing for the construction of a functional connectivity map (Ito et al. 2014; Miccoli et al. 2019; Schulte et al. 2021). By using HD MEA, we are able to detect changes in network activity both in primary neuron cultures and in brain slices.

This study examines the physiological significance of NLF from 5XFAD animals in altering neuronal physiology and connectivity. Using the microfluidic sorting technique, we are able to collect morphologically intact exosomes from the NLF which are applied to primary neuronal cultures and organotypic hippocampal slice cultures (OHSC). The functional effects of exosomes on neuronal firing and network are meticulously characterized by an HD MEA system. As a result, connectivity analysis revealed a progressive decrease in network efficiency in neuronal cultures under the exosomes from 5XFAD NLF and Aβ42 oligomer over time. In addition, local field potential (LFP) recordings in OHSC also demonstrated aberrant elevation in the same conditions. It is plausible that these alterations of neural networks in response to 5XFAD NLF could lead to disseminating the neurodegenerative potential of substrates contained within exosomes.

## Results

### Characterization of exosomes isolated from nasal lavage fluid of 5XFAD mice using FAST method

To examine the role of exosomes from NLF of 5XFAD animals, the flow amplification separation technology or FAST was applied to collect intact exosomes from miniscule amounts of nasal fluid in high yield and purity (Shin et al. 2017) (Figure 1B). Under the consistent temperature control, we obtained the size distribution of the particles isolated from NLF through nanoparticle tracking analysis or NTA (Figure 1D). Compared to the control, the overall distribution pattern of 5XFAD represented higher peaks in particle concentration. The concentration of 35∼205 nm particles isolated from the nasal fluid of 5XFAD mice was about 8.5 × 10^8^ particles/ml, and the value was about 3.0 folds higher than the control (Figure 1E). According to the transmission electron microscopy (TEM) image, majority of collected exosomes fell into size in rage of 30 ∼ 150 nm diameter (Figure 1F). Apparently, the density of exosomes in the same volume loaded onto the grid was found higher in 5XFAD mice that was confirmed with 3.2 folds increase of CD63 exosome marker expression level (Figure 1G and H). Overall, the secretion of NLF-derived exosomes was enhanced in Alzheimer’s animal model similar to other body fluids (Rajendran et al. 2006; Saeedi et al. 2019).

**Figure 1.**
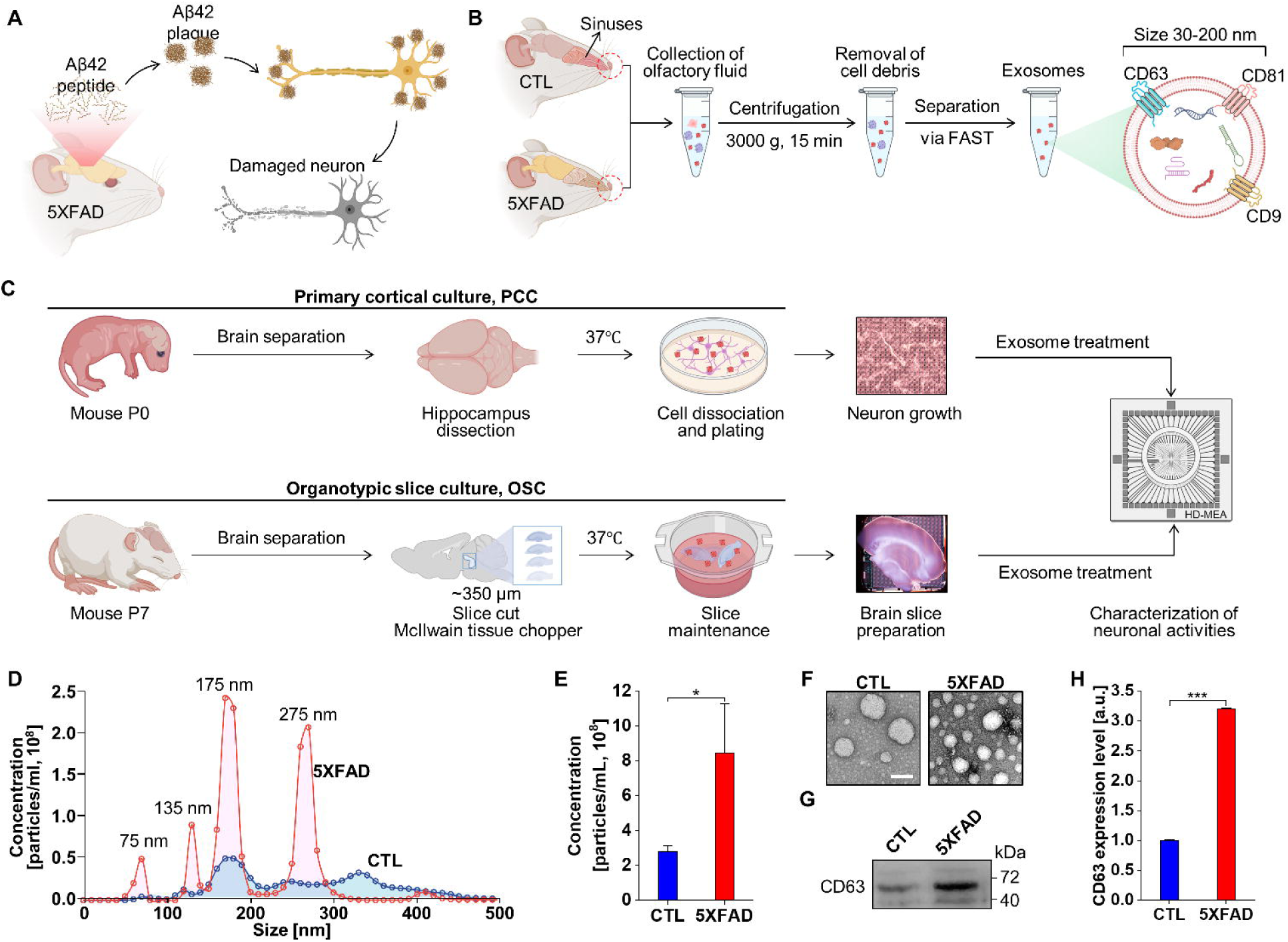
The workflow of exosome preparation from NLF and its quantification following functional tests in HD MEA. (A) Schematic drawing of Aβ42 accumulation in the brain of Alzheimer’s model animal. (B) Flow amplification separation technology, FAST method to collect intact exosomes from NLF of 5XFAD and control animals. Tetraspanins expression in the vesicle membrane such as CD63, CD81, and CD9 as markers of exosomes (C) Experimental flow of functional tests of exosome in the primary cortical culture and the organotypic slice culture with the successive HD MEA recording (D) Size distribution of particles isolated from NLF through nanoparticle tracking analysis. NLF exosomes from 5XFAD in red, control in blue. (E) Particle concentration in the range of 35∼205 nm. (F) TEM images in the grids of same volume loading with 5XFAD and control NLF-derived exosomes. Scale bar = 100 nm. (G) Western blot of the exosome surface marker CD63. (H) Protein expression level of exosome surface marker CD63. The values are means ± SEM from three independent experiments. **p<.01, *p<.05; unpaired, two-tailed t test with Welch’s correction.

### Alteration of neuronal excitability and network connectivity in the primary cortical neuron cultures induced by Aβ42 oligomers and 5XFAD NLF-derived exosomes

During the development stage of Alzheimer’s symptom, neuronal excitability has often been observed in vitro cell studies with pathologic Aβ42 oligomer treatment as well as fMRI imaging in MCI patients. In this study, primary cortical neurons were recorded for two weeks by HD MEA in order to measure the neuronal excitability induced by Aβ42 oligomer. In addition, 5XFAD NLF-derived exosomes were also tested under the same conditions to compare the neuronal excitability. The basic topological properties of neural cultures were assessed at days *in vitro* (DIV) 7, 10, and 13 based on the number of spikes (NOS), interspike interval (ISI) and mean firing rate (MFR) for each group. During the culture ageing progress, NOS and MFR tended to increase in all groups (Figure 2A, C and D). Among them, the Aβ42 treatment group displayed the strongest level of activity. Although the NOS in the control group was slightly higher than that in the 5XFAD group, the MFR was in the opposite case. ISI as an indicative of the time interval between two consecutive spikes, tended to decrease as the culture aged implying the increased readiness in response to an input (Figure 2E). An analysis of bursts in a network provides useful information about the maturity of the network. As a network matures, its neurons become more capable of receiving multiple inputs, both excitatory and inhibitory, more frequently, thereby extending the duration of bursts. For these functions to be maintained for a longer period of time, robust ion pumps and channel machinery are required to maintain ionic homeostasis and prevent runaway or rundown of connectivity (Boehler et al. 2012; Ellender et al. 2010). As the cultures aged, the burst duration increased monotonically in all the groups (Figure 2B and F). The mean number of spikes per burst remained consistent across different groups and DIVs suggesting stable network mechanisms underlying individual burst events (Figure 2G). The burst frequency, however, was apparently increased in Aβ42 and 5XFAD NLF groups in older cultures indicating more recurrent generation of neuronal activation in those detrimental conditions (Figure 2H). In the burst patterns in neuronal networks, it was also evident that the network burst duration in interquartile range showed steady increase in Aβ42 and 5XFAD NLF groups compared to the indistinguishable change in control (Figure 2I).

**Figure 2.**
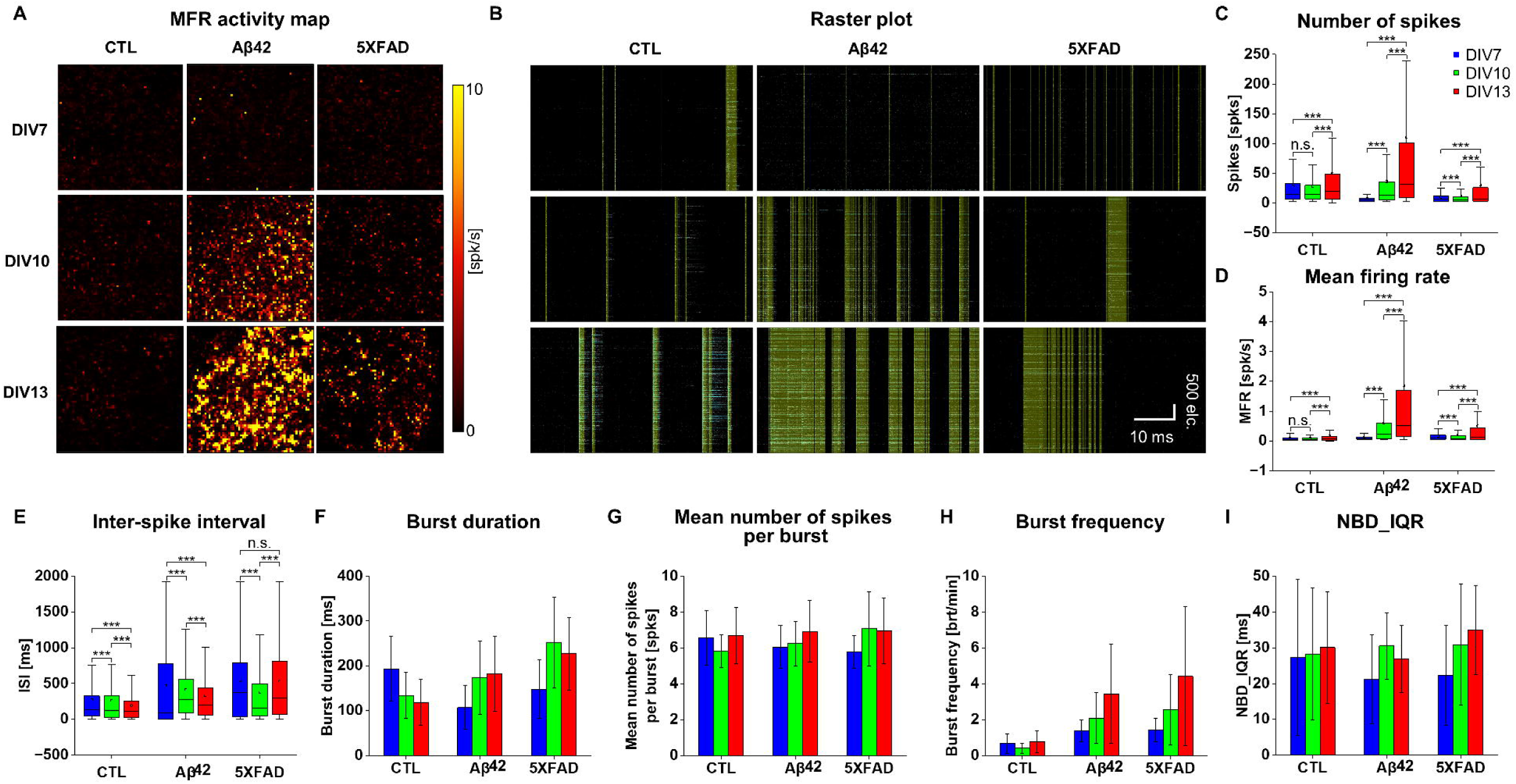
Topological properties of neuronal cultures on HD MEA recording with culture development. (A) MFR Activity map in spontaneous neuronal activation during 60 seconds recordings. Control, Aβ42, and 5XFAD NLF treatment groups in DIV 7, 10, 13. The intensity scale range from 0 to 10 spikes per second. (B) Raster plot of neuronal spiking from 4096 electrodes (y axis) for 60 seconds recording (x axis). Mustard color represents for spike network bursts, and blue for spike bursts. (C) Box plot of number of total spikes for 60 seconds recordings in the individual culture chips. Blue for DIV 7, green for DIV 10, and red for DIV 13. (D) Box plot of mean firing rate (MFR) in spikes per second. (E) Box plot of inter-spike interval (ISI) in millisecond. (F) Average of burst duration with error bar in millisecond. (G) Average of mean number of spikes per burst in the raster plot with error bar (H) Average of burst frequency in bursts per minute with error bar (I) Average of network burst in interquartile range in millisecond. In all the box plots above, lower quartile as the borderline of the box nearest to zero expresses the 25th percentile, whereas upper quartile as the borderline of the box farthest from zero indicates the 75th percentile. Error bars show SEM. ***p□<□.005, **p□<□.01, *p□<□.05; unpaired, two-tailed t test with Welch’s correction; n.s. not significant.

### Differentiating features of neuronal networks in 5XFAD NLF treated neurons

To investigate network properties, it is prerequisite to apply reliable network parameters to describe its essential characteristics (Mossink et al. 2021). Among the many parameters, clustering coefficients (CC), node degrees (ND), and path lengths (PL) were recruited due to the credibility and essentiality. First of all, CC is primarily a measure of the tendency for neurons in a network to form clusters. In that sense, the clustering of the network gradually increased in all the groups (Figure 3B) albeit the sparser connections in the connectivity map, particularly from DIV10 to 13 (Figure 3A). We presented path length (PL) and node degree (ND) as average values and distribution histograms for each node (Figure 3C-F).

**Figure 3.**
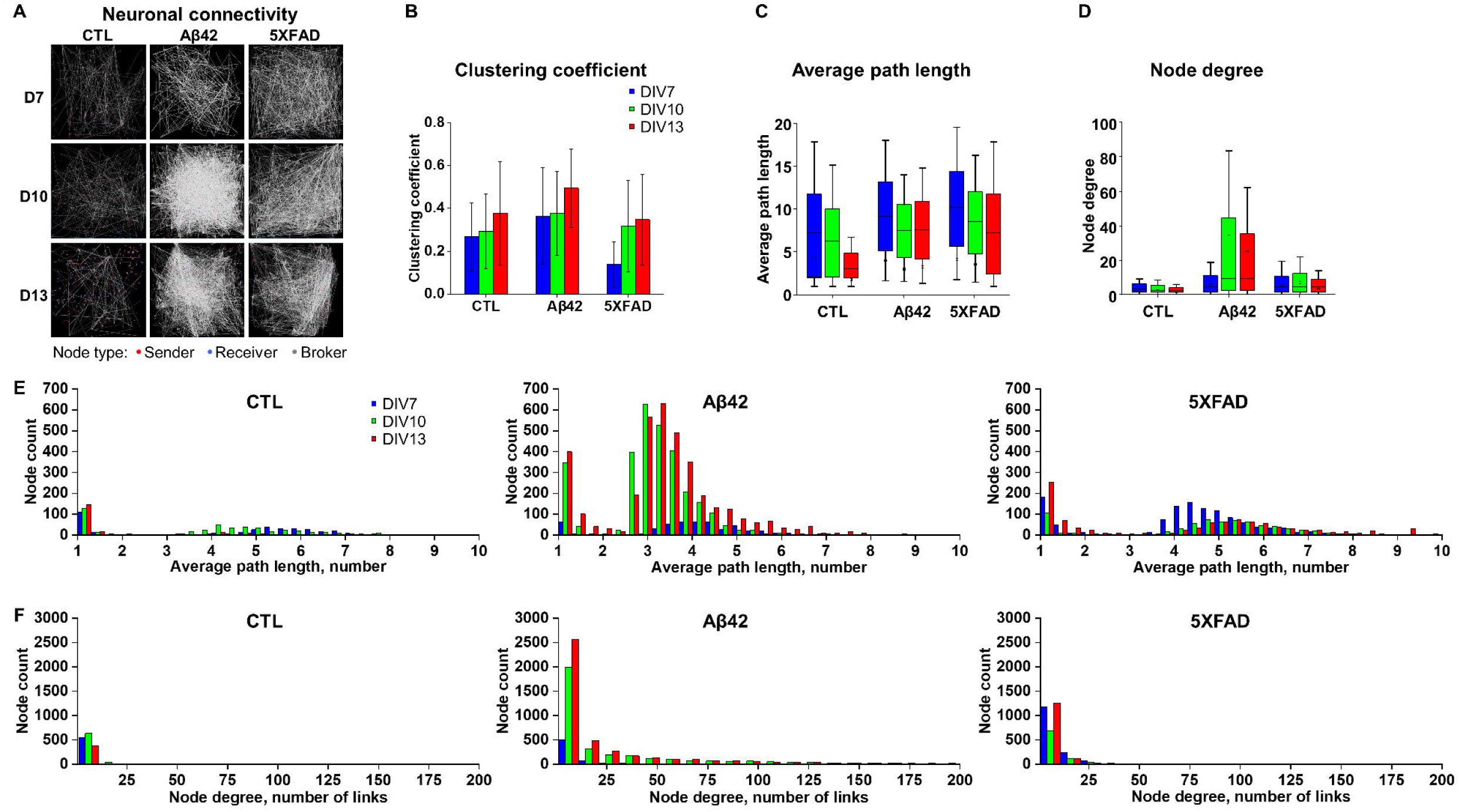
Characteristics of neuronal connectivity in networks. (A) neuronal connectivity map in spontaneous neuronal activation during 60 seconds recordings. Control, Aβ42, and 5XFAD NLF treatment groups in DIV 7, 10, 13. Red dot represents for a node of sender, blue for receiver, and gray for broker. White line describes connection between nodes. (B) Average of clustering coefficients from neuronal connections for 60 seconds recordings in the individual culture chips. Blue for DIV 7, green for DIV 10, and red for DIV 13. (C) Box plot of average path length in number of links (D) Box plot of node degrees (E) Distribution histogram of node counts to average path length in number. (F) Distribution histogram of node counts to node degree in number of links.

PL measures the number of steps involved in transmitting signals from one node to another. In the control culture, the PL value steadily declined (Figure 3C Left), suggesting that neuronal communication had become more efficient over time. Given the increasing CC, this phenomenon can be due to keeping more long-distance shortcut connections as well as local ones (Watts and Strogatz 1998). In contrast, the PLs in the Aβ42 and 5XFAD NLF groups retained the high means and wider deviations as reflected in the boxplots (Figure 3C Middle, Right). The histograms for the PL distribution also revealed that the tails in the PL distribution became fatter in older DIVs in comparison with the control group (Figure 3E and Figure S1A). Those data suggested that the networks of the Aβ42 and 5XFAD groups added or kept more local, clustering connections over time, which increased CCs, but not enough long-distance shortcuts, leading to the degraded network efficiency, compared to the control.

ND, representing the number of connections per node, provided further insights for the network changes over time, contrasting between the control group and others. In the control group, ND gradually declined, suggesting the network-wide pruning of connections. However, in the Aβ42 group, there was a substantial leap in the number of connections from DIV7 to 10 followed by a slight decrease in DIV13 (Figure 3A and D) while 5XFAD also showed similar tendency in lesser degree. The ND distribution histogram reflected the quantitative changes across DIVs (Figure 3F and S1 B).

In summary, the control group network evolved in time by losing connections while it still developed higher CC and lower PL, developing a more efficient, ‘small world’-like structure(Watts and Strogatz 1998). However, in both the Aβ42 and 5XFAD NLF groups, the networks added (from DIV 7 to 10) and lost connections (from DIV10 to 13) while the latter did not improve efficiency much, likely due to a broken balance between the local connections and long-distance shortcuts. This result demonstrates that 5XFAD NLF caused qualitatively the same pathological changes in the network activity patterns as Aβ42.

### The effects of Aβ42 oligomers and 5XFAD NLF-derived exosomes on the LFP property and oscillation in organotypic hippocampal slice cultures (OHSC)

Neuronal culture has limitations in its artificial construction, as opposed to the intrinsic circuitry formation in the brain. To investigate the network properties of neurons in a more natural environment, we tested OHSC incubated with Aβ42 oligomer and 5XFAD NLF-derived exosomes with HD MEA. Under 4AP stimulation, LFP signals in slices were recorded after 12 days of incubation with each of the treatments. Typical LFP traces for the dentate gyrus (DG), cornu ammonis (CA) 1 and 2 have been presented in Figure 4A, overlaid on the MEA recording. As in neuronal cultures, internal connections within slices were represented in OHSC overlapped on top of MFR responses (light blue lines, Figure 4A). After low-pass filtering (< 200 Hz), certain types of oscillations were evident in DG, CA1, and CA3 of every OHSC (Figure 4A). LFP analysis revealed that Aβ42 had the highest average amplitude following 5XFAD NLF and control, but that 5XFAD NLF still had the widest deviation range (Figure 4B). Further, 5XFAD NLF maintained both the highest average value and a wider deviation in LFP rate and duration (Figure 4C and D). In spite of differences in LFP characteristics, the energy levels in all slices remained the same, suggesting that activity dynamics were consistent across the test samples (Figure 4E).

**Figure 4.**
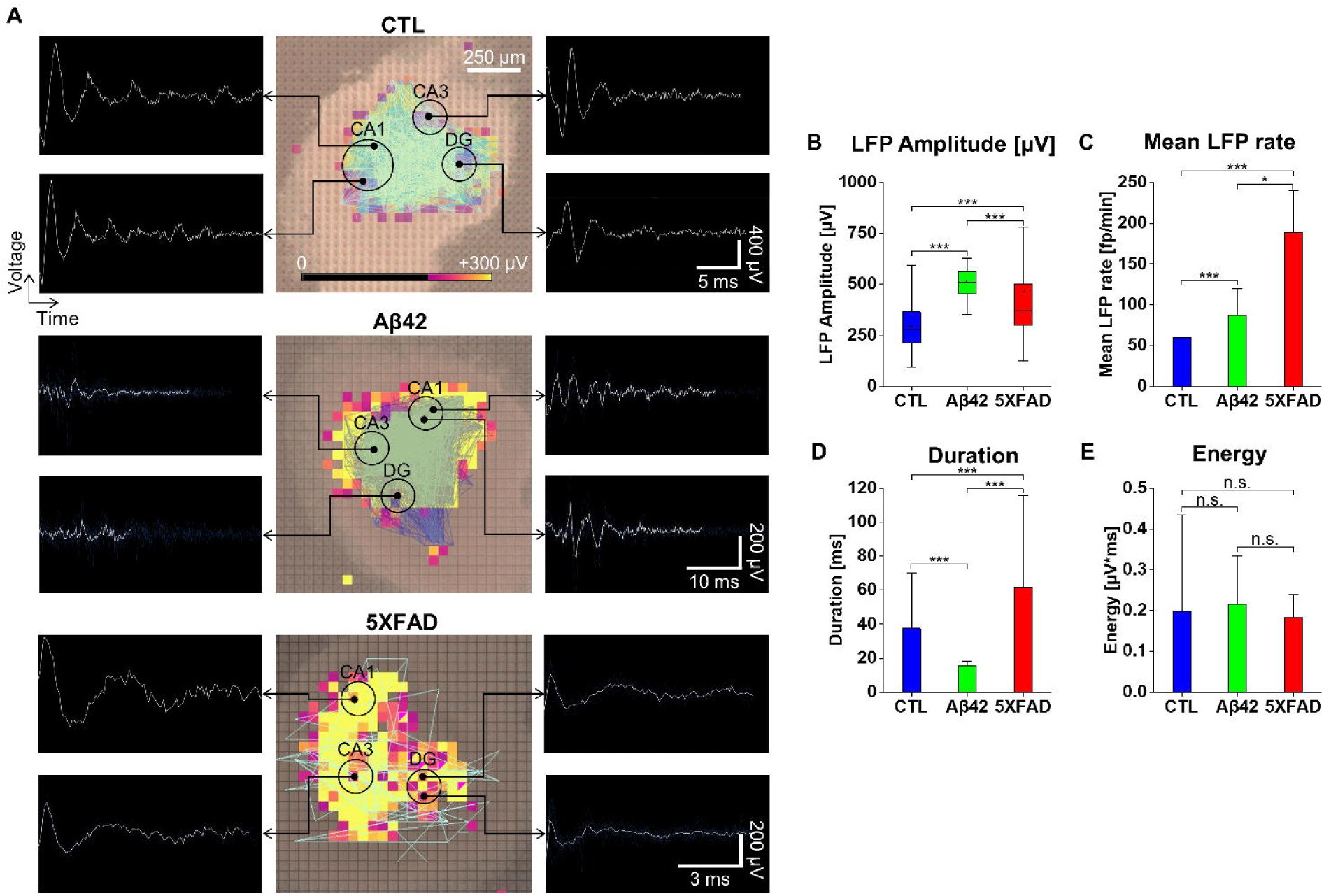
MEA record of LFP responses in organotypic hippocampal slices (OHCS) (A) Overlay of LFP activities on the recording OHCS with distinctive waveforms in DG, CA1, and 2 regions. Control, Aβ42, and 5XFAD NLF treatment groups after incubation of 12 days. Intensity of LFP activity in the color scale from 0 to 300 µV. The sky blue line represents for connections within the slices. LFP waveforms after lowpass filter (< 200Hz). (B) Box plot of LFP amplitude in micro voltage (C) Average of mean LFP rate in number of LFP per minute with error bar (D) The average duration of LFP in millisecond with error bar (E) Average of mean LFP energy in charge unit of measurement for the area under a voltage-time curve. Error bars show SEM. ***p□<□.005, **p□<□.01, *p□<□.05; unpaired, two-tailed t test with Welch’s correction; n.s. not significant.

### Current source density (CSD) analysis to localize LFP distribution in the organotypic hippocampal slice cultures (OHSC)

Even though LFP analysis can deliver the characteristic neural activity in brain slice, the vulnerability of far-field effect to volume conduction is the intrinsic limitation of LFP in terms of signal transmission in the recording slices. To overcome the caveat, a second spatial derivative of LFP in other word, current source density (CSD) can be applied to estimate the distribution and propagation of LFP in high accuracy (Nicholson 1973; Nicholson and Freeman 1975). A comparison of the mean CSDs across the groups indicates that 5XFAD NLF group has the highest level implying significantly larger electrical current sources than sinks on recording slices (Figure 5A). Although the mean value of the Aβ42 group was lower than that of the 5XFAD NLF, the deviation was wider, suggesting that the data for individual slice are complex and spread in between the control and 5XFAD NLF cases. The propagation of LFP converted to CSD was recreated into time series motion with sinks and sources (Mov. 1 to 6). The representative images were captured with snap shots during the time of activity spread (Figure 5B,C). Parsing of the CSD signals separated the sink and source waveforms where the level and frequency of ridges and furrows of wave spread was manifest (Figure S2 A to C). Apparently, 5XFAD NLF showed the most volatile dynamics, followed by that of the Aβ42-treated OHSC.

**Figure 5.**
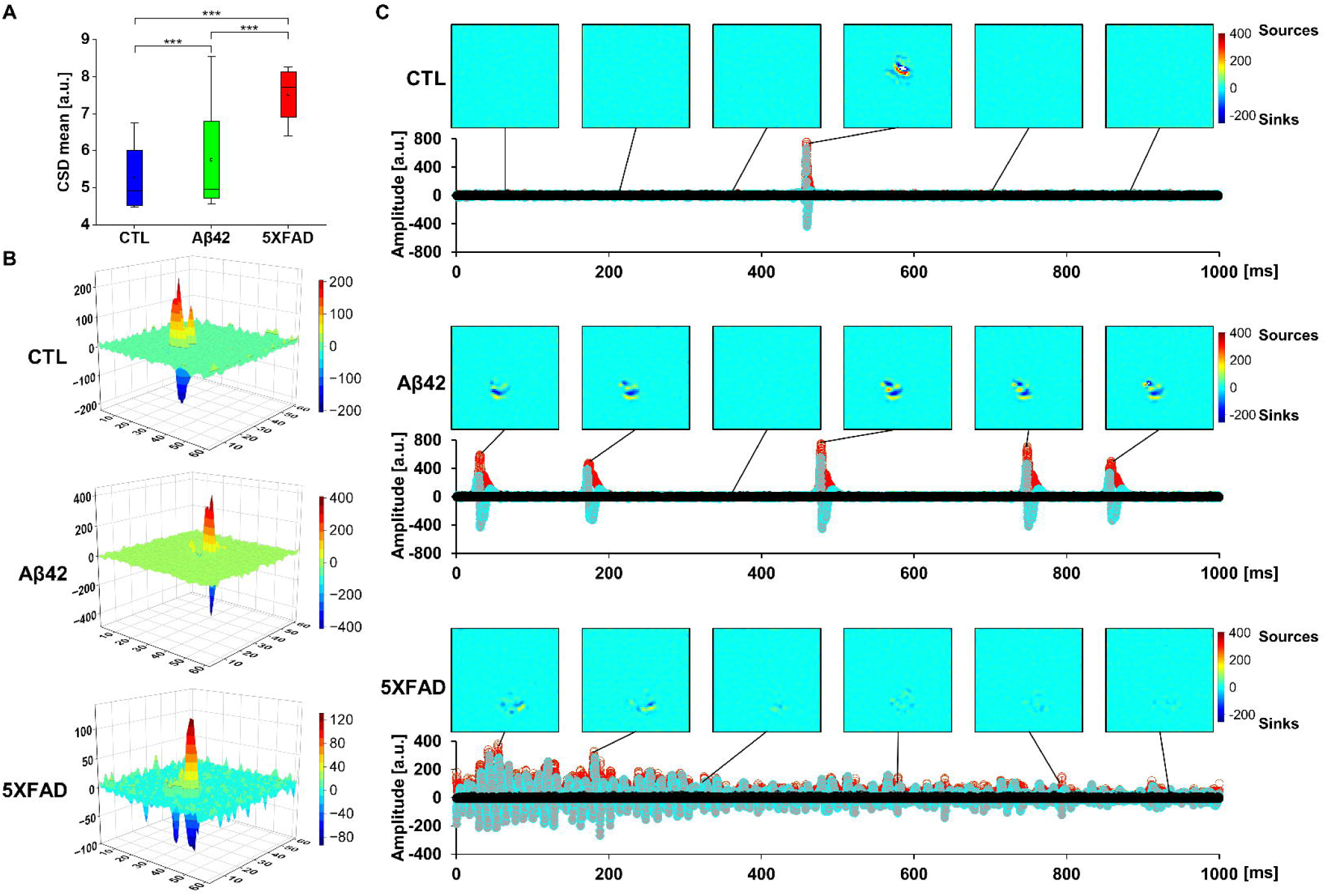
CSD propagation in organotypic hippocampal slices (OHCS) (A) Box plot of CSD mean in arbitrary unit. (B) Snap shot of CSD amplitude changes in 3D graph captured from MOVIE 1-3 of respective CSD propagations. Blue for sinks and red for sources in scale bar. (C) Distribution of CSD incidences over time in amplitude. The upper images are captured from MOVIE 1-3 of respective CSD propagations. Blue for sinks and red for sources in scale bar.

## Discussion

Exosome research has experienced an explosive growth in recent years, particularly in perspective of cellular mediation, which is useful for AD studies as well. Previously, it has been demonstrated that Aβ42 pathology can propagate in a similar manner to the transmission of prion disease (Shankar et al. 2008). The actual mechanism, however, by which Aβ42 is transferred between cells was still elusive. The presence of exosomes containing amyloid precursor peptide (APP) and Aβ42 has been implicated in AD pathogenesis, potentially increasing the spread of pathologic agents (Howitt and Hill 2016; Sharples et al. 2008; Yuyama et al. 2012). Direct evidence was found in post-mortem brain samples of AD patients from which pathologic Aβ42 plaque was localized in the dissociated exosomes (Sardar Sinha et al. 2018). In the study, the contagious property of the exosomes was confirmed in neuronal cultures that propagated exosomes upon engulfing them and then infected neighboring neurons. Thus, we wanted to address the effect of AD brain-derived exosomes in neuronal activity modulation. Rather than dissociating exosomes from brain samples, nasal lavage fluid (NLF) was prepared because it can offer a convenient method to collect nasal wash in large quantities and on a noninvasive manner in clinical settings. We hypothesized that exosomes in NLF potentially contain AD-related pathologic molecules considering the tight correlation between olfactory dysfunction and dementia symptom (Attems and Jellinger 2006; Bathini et al. 2019; Doty 2017; Marin et al. 2018). The microfluidic sorting technology allowed us to obtain intact exosomes at the femtomolar level after extracting clean nasal wash from mice using the fast nasal irrigation method (Cho et al. 2012; Shin et al. 2017). In spite of this, characterization of the internal contents was confronted with technical obstacles due to the insignificant amount of exosomes collected from nasal washes. For proteomics or RNA sequencing analysis, the collection of a sufficient volume of nasal wash is a prerequisite for running the analysis procedures on an adequate number of subjects. Additionally, the construction of a library set with sophisticated analysis programs is another challenge whenever novel samples are being tested for the first time. Due to the fact that such a study is beyond the scope of the current design of experiment, it will be kept as a follow-up assignment.

Neuronal hyperactivity has been implicated in epileptic seizures in both humans and AD model animals as an unique feature of early stages of AD (Palop et al. 2007; Verret et al. 2012; Vossel et al. 2013). The electrophysiological results in this study with soluble Aβ42 oligomer treatment on neuronal cell cultures were consistent with those reported previously regarding hyperexcitability of neurons (Busche et al. 2012; Busche et al. 2008; Muller et al. 2021; Zott et al. 2019). Surprisingly, 5XFAD NLF exosome treatment also enhanced neuronal activity as well, consistent with a report on hippocampal hyperactivity conducted by whole-cell patch clamp in 5XFAD brain slices (Andersen et al. 2021; Li et al. 2021). According to the research, it is attributed to attenuation of inhibitory postsynaptic current and metabolic dysfunction between neurons and astrocytes in 5XFAD hippocampal neurons. The effect of exosomes on functional signaling networks was reported in MECP2-knockdown human primary neural cultures where induction of neurogenesis, synaptogenesis, and circuit connectivity was observed in hiPSC-derived exosome treatment (Sharma et al. 2019).

Due to the exceptional integrity of HD MEA recordings from 4096 electrodes in real time, it has been possible to systematically determine the connectivity between neuronal networks. The gradual increase of clustering coefficient values in aging cultures indicates that network stability can also be developed over time in neurodegenerative conditions such as Aβ42 oligomer and 5XFAD NLF exosome treatment. According to the quantitative dimension of network, such a pathological environment stimulates the formation of clusters within neuronal cultures that are relevant to neuronal hyperexcitability. Apart from the superficial observation of the network, the qualitative analysis calculated from the path lengths between nodes revealed that disintegration of network efficiency occurs during a longer incubation period with 5XFAD NLF exosomes at the same rate as it does with Aβ42.

A longitudinal study of soluble Aβ42 oligomers and 5XFAD NLF exosomes on OHSC was conducted to determine their neuromodulatory effect in mouse hippocampal slices. There were also increases in LFP responses in Aβ42 oligomers and 5XFAD NLF exosomes on OHSC, which mirrored the increase in neuronal activity observed in the dissociated cultures. It was observed in the neuronal culture that Aβ42 oligomer treatment exhibited the strongest topological activation characteristics as well as connectivity. However, the LFP property was higher in 5XFAD NLF exosomes than Aβ42 oligomer treatment, which was also reflected in the CSD analysis. In this case, it is likely that the intrinsic structure of hippocampal slices consists of neurons and glial cells interspersed within a specific circuit. An altered current conduction in a hippocampal slice may result from the interaction of substances in NLF exosomes with diverse cells in the circuitry.

As AD research has mainly focused on elucidating neurobiological pathophysiology, very little knowledge has been revealed regarding neuronal electrophysiology and its network properties of AD brain. This limitation is a result of the technological impediment of being unable to observe the whole picture of individual and inter-neuronal function simultaneously. By employing HD MEA recording, we were able to demonstrate the functional effect of exosomes derived from NLF in AD model animals. The exosomes enhance neuronal activity and network connectivity in a similar manner to soluble Aβ42 oligomers, which is often present in the early stages of Alzheimer’s disease. This study provides an opportunity to apply exosomes from olfactory fluid as a biomarker of Alzheimer’s disease in clinical examinations as well as in biological research.

## Materials and Methods

### Animals

B6SJL-Tg (APPSwFlLon,PSEN1*M146L*L286V) (5XFAD) mice were purchased from The Jackson Laboratory (MMRRC Stock No: 34840-JAX) and experimental procedures were performed according to protocols approved by the Institutional Animal Care and Use Committee (IACUC) of KPCLab (approved number: P171011) and ⓒMEDIFRON DBT Inc. (approved number: Medifron 2017-1). C57BL/6 mice were obtained from OrientBio Inc. and compliance with relevant ethical regulations and animal procedures were reviewed and approved by Seoul National University Hospital IACUC (approved number: 16-0043-c1a0).

### Nasal lavage fluid extraction

The procedure of nasal lavage fluid (NLF) extraction was followed to the previous method (Cho et al. 2012). After anesthesia, the left ventricle of mice was cannulated and blood was cleared by perfusion with cold PBS. Using scissor dissection, the upper airway, including the palatopharyngeal region, was separated, and the mouse head was separated at the larynx level of the upper airway. Two consecutive volumes of 350 µl of PBS were instilled through the pharyngeal opening into the choana. NLF fluids were centrifuged, and supernatants were stored at 80°C until assayed.

### Exosome purification

Nasal-derived exosome isolation was performed as described previously (Shin et al. 2017), with minor modifications. In brief, nasal lavage fluid harvested from the mouse model was filtered through a 0.2-µm syringe filter (Sartorius, Goettingen, Germany) to remove aggregates. The nasal fluid, including extracellular vesicles, was carefully collected and kept on ice before performing exosome separation using the FAST. To isolate exosomes, purification buffer was filtered through a 0.2-µm syringe filter. The flow ratio of sample:buffer:magnification was set at 5:95:75. Exosome-sized particles were separated from the other particles and all samples were maintained at 4°C during exosome purification.

### Nanoparticle tracking analysis

NTA was performed using an LM10 (NanoSight, Salisbury, UK) instrument. Exosomes separated from miniscule amounts of nasal lavage fluid were diluted with filtered phosphate-buffered saline (PBS) to examine 20 particles per frame and gently injected into the laser chamber. Each exosome sample was subjected to a red laser (642 nm) three times for 1 min each; the detection threshold was set to 5 to allow the detection of nanosized particles. The data were analyzed using NTA software (ver. 3.1; NanoSight). All experiments were conducted at room temperature.

### Western blotting

CD63 expression was quantitatively analyzed using Western blotting. Purified exosomes were mixed with 5% sodium dodecyl sulfate (SDS) sample buffer (Tech &Innovation, Korea) and sliced tissues were lysed and homogenized in radioimmunoprecipitation assay buffer (Tech &Innovation, Korea) containing protease inhibitors. The protein concentrations of the separated solutions were measured using the Bradford assay (Bio-Rad, Hercules, CA, USA). The samples were heated for 10 min at 97°C. Next, 15 µg of protein from each sample were subjected to SDS-polyacrylamide gel electrophoresis (PAGE) (12%) and transferred to polyvinylidene difluoride (PVDF) membranes (Bio-Rad) for 90 min. Each membrane was blocked with 5% skim milk (BD) in TBST buffer (25 mM Tris, 190 mM NaCl, and 0.05% Tween 20, pH 7.5) for 1 h at room temperature, followed by incubation with primary antibodies (anti-CD63, 1:300; Novus Biologicals, Centennial, CO, USA) at 4°C overnight. After five 20-min washes with TBST, each membrane was washed three times in Tween-20 and incubated with goat anti-mouse IgG for 2 h. Bands were visualized using an enhanced chemiluminescence system; the intensity of the blots was quantified with ImageJ software.

### Transmission electron microscopy

Separated exosomes were diluted and fixed with 2% glutaraldehyde overnight at 4°C. The mixture of exosomes and fixation solution was then diluted 10-fold with PBS for electron microscopic observation. Briefly, 5 µL of each sample was plated onto a glow discharged carbon-coated grid (Harrick Plasma, Ithaca, NY, USA), which was immediately negatively stained using 1% uranyl acetate (ref2). The exosome samples on the grids were observed under a Tecnai 10 transmission electron microscope (FEI, Hillsboro, OR, USA) operated at 100 kV. Images were acquired with a 2k × 2k UltraScan CCD camera (Gatan, Pleasanton, CA, USA).

### Aβ42 oligomer preparation

The peptide corresponding to human Aβ42 (Anaspec, AS-64129-1, 1 mg) was dissolved in 100 μL of DMSO by vortexing for 30 minutes at room temperature, and then the solution was added to 900 μL of PBS for incubation at 4°C for 24 hours.

### Electrophysiology

### Primary neuron culture

Dissection medium Neurobasal Media (NBM) consisted of 45 ml Neurobasal Medium A, 1 ml B27 (50 X), 0.5 mM Glutamine sol, 25 μM Glutamate, 5 ml Horse serum, 500 μl penicillin/streptomycin. And culture medium consisted of 50 ml Neurobasal Medium A, 1 ml B27, 0.5 mM Glutamine sol, 500 μl penicillin/streptomycin, 50 μl HEPES. The Biochip chamber (3Brain, Arena) was cleaned, filled with 70 % ethanol for 20-30 minutes, rinsed with autoclaved DDW for 3-4 times and dried in the clean bench overnight with NBM. On the day after, 30-90 μl filtered PDLO which dissolved in borate buffer on the active surface of the Biochip was added and placed overnight in the incubator. The Biochip was washed with autoclaved DDW for 3 times before cell seeding. Primary cortical and hippocampal neuron culture were prepared from postnatal 0-day mouse pups. Pups were decapitated with sterilized scissors and the whole brain was removed. The removed brain was chilled in cold neurobasal medium with papain 0.003 g/ml solution at 4°C in a 35 mm diameter dish. Surrounding meninges and excess white matter were pulled out under the microscope (Inverted microscope, Nikon, Japan) in the same medium to a second dish at 4°C. The cortex and hippocampus parts were isolated from other parts of the brain, washed with NBM and papain solution, and minced into small pieces. The minced tissues were transferred into a 15 ml tube and incubated for 30 min in the 37°C water bath. After, the tube was inverted gently every 5 min to be mixed. The tissues were washed with HBSS twice, after being settled down, the cortex and hippocampi tissues were transferred into prewarmed NBM and triturated for 20-30 times using a fire-polished Pasteur pipette. The number of cells was counted and 30-90 µl drops of the cells were plated in the Biochip, which contains ∼1000-1500 cells/µl (incubated in 37°C in 5 % CO_2_). From day 2 of the culture, the whole medium was replaced with a fresh feeding medium every 3 days. 10 µM Aβ42 oligomers was treated at DIV7, 10, and 13. 5XFAD NLF exosomes were incubated from DIV7 until DIV13.

### Organotypic hippocampal slice culture

Dissection medium consisted of 40 ml Hibernate A, 0.5 mM L-Glutamine, 10 ml Horse serum. Growth medium 1 consisted of 40 ml Neurobasal A, 20% Horse serum, 400 μl penicillin/streptomycin solution. Growth medium 2 consisted of 40 ml Neurobasal A, 2% B27 supplement, 400 μl penicillin/streptomycin solution. The organotypic hippocampal slice culture was prepared from postnatal 7-day mouse pups. Decapitation was performed under cervical dislocation and the entire brain was removed with forceps and chilled in Hibernate A medium for 10 seconds at 4°C. After that, the excess white matter and meninges surrounding the brain were removed under the microscope (Inverted microscope, Nikon, Japan) carefully and the hippocampus was dissected with a spatula. The dissected hippocampus was placed on the tissue chopper instrument (Stoelting tissue slicer 51425, SA) which has filter paper coverage and sliced transversally to 300 µm. The freshly cut sections were collected into the cold dissection medium and separated from each other by spatula. From the best slices, up to 4 slices were transferred onto the cold culture membrane (Millicell membrane, 0.4 μm) which then placed into the prewarmed 6-well plate with Growth medium-1. From day 2, the medium was changed with Growth medium-2 and the whole medium was replaced every 3 days. 10 µM Aβ42 oligomers and 5XFAD NLF exosomes were treated at DIV7 until DIV13.

### Neuronal spike and LFP recording with High density Multielectrode array (HD-MEA)

High density Multielectrode array (HD-MEA) recording with 4096 electrodes in CMOS Biochip (BiocamX, 3Brain GmbH, Switzerland) was conducted at sampling rate of 9 kHz. The active electrode which is 21 μm × 21 μm in size and 42 μm in pitch is implanted in the array with 64 × 64 grid (2.67 × 2.67 mm^2^) centered in a working area (6 × 6 mm^2^). The brain slice was positioned at the center of the Biochip under the square shape platinum net anchor to prevent it from displacement by the perfusion flux. In order to overlay the real image of the slice at the site of recording, a stereomicroscope with transmitted illumination was settled over the slice with 20 × magnification (Nikon, SMZ745T, Japan). The neuronal activities in cortex slices were recorded for approximately 35 min with and without additional chemicals. In order to generate spontaneous epileptic like discharges, the slices were perfused with Kv1 channel blocker 4-Aminopyridine (4AP) 250 μM (Sigma-Aldrich, USA). Spontaneous response was recorded for the first 5 min following the oxygenated 4AP for up to 15 min. All recordings were conducted by Brainwave software (3Brain GmbH,Switzerland).

### HD MEA data analysis

Raw data was filtered with a IIR low pass filter (cut off at 200 Hz, order 5) before LFP detection. To identify LFP events, a standard hard double threshold algorithm was used (upper threshold 40 µV, lower threshold -40 µV). When the signal overcame one of the two thresholds, an LFP was detected. To determine the duration of the LFP, the energy of the signal was calculated on a sliding window of 50 ms moving forward and backward around the peak until the energy was 1.5 times higher than the energy calculated on the noise. A refractory period (i.e. the minimum distance between two consecutive LFPs on an electrode) of 50 ms was set by the operator. An electrode was considered active if the LFP rate was at least 0.05 event/sec. After detection, statistics on the LFP features were extracted by averaging the parameter on each electrode along the recording. Then distribution of the feature was calculated by grouping together electrodes belonging to the same anatomical area. To identify the different areas with respect to the electrodes position, an image of the recorded slice was superimposed to the map of the electrodes grid, allowing manual selection of electrodes for each area of interest. LFP event generally involved most of the area of interest, so to get rid of spurious false positive detected events, an automated cleaning procedure was used before feature extraction. Detected LFPs were considered valid only if they were occurring simultaneously on at least the 40% of the total electrodes of an area within a time window of 300 ms. All the parameters from spike detection and LFP recording were calculated with brainwave software (3Brain).

### Current-source density (CSD) analysis

In the two-dimensional CSD analysis, we first preprocessed the data in two steps: first, at each time point, we identified the saturated electrodes by simple thresholding and inpainted missing signals by linear interpolation with the data from surrounding regions. We monitored how much space the saturated electrodes occupied and checked whether this procedure caused any noticeable artifacts. Then, we smoothed the data spatially by a gaussian kernel with σ = *d_pitch_*. From this, the current source is estimated by applying the modified two-dimensional Laplacian (Ortiz et al. 2018)

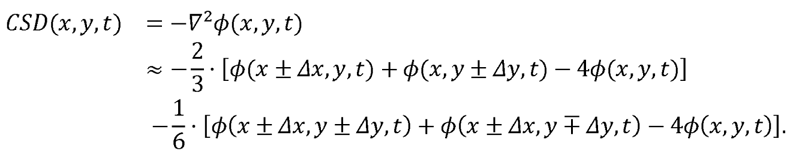

From this, we isolated CSD within the region of interest (ROI) and time window of interest (TOI), computed in the following way: We first estimated the dynamic amplitudes of LFP signals *A*(*x*,*y*,*t*) by applying Hilbert transformation (MATLAB function *hilbert*) and computing their absolute values. Then, for each (*x*, *y*), we computed the amplitude variability, ξ(*x*,*y*) = STD[*A*(*x*,*y*,*t*)]*_t_*, and the ROI was selected by a criterion, ξ(*x*,*y*) > Median[ξ(*x*,*y*)] + 2.326 STD[ξ(*x*,*y*)]. For each (*x*, *y*) in the ROI, a TOI was selected by *A*(*x*,*y*,*t*) > μ_noise_ + 2.326 √ 2 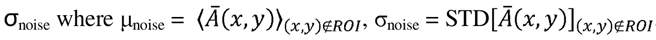, and *Ā(x,y)* is the temporal average of *A*(*x*,*y*,*t*). Then, the average rectified CSD (rCSD) is computed by averaging the absolute value of CSD,|CSD(*x,y,t*)|, within the ROI and TOI.

All CSD analysis was done by custom scripts in MATLAB 2018a (Mathworks Inc., MA), which will be available upon request.

## Supporting information

MOV1

MOV2

---

**Supplementary Figure 1.**
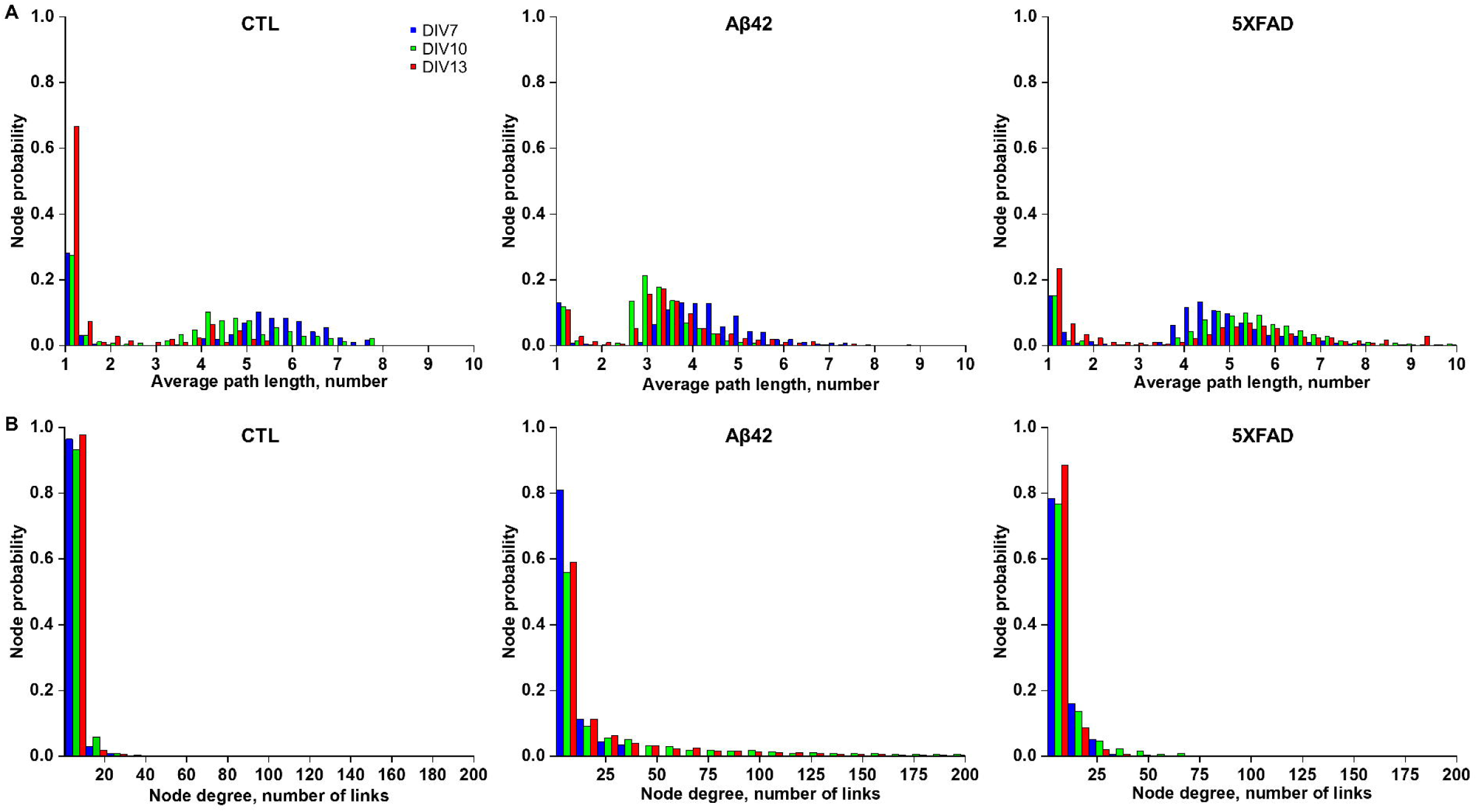
(A) Distribution histogram of node probability to average path length in number. Blue for DIV 7, green for DIV 10, and red for DIV 13. (B) Distribution histogram of node probability to node degree in number of links. Blue for DIV 7, green for DIV 10, and red for DIV 13.

**Supplementary Figure 2.**
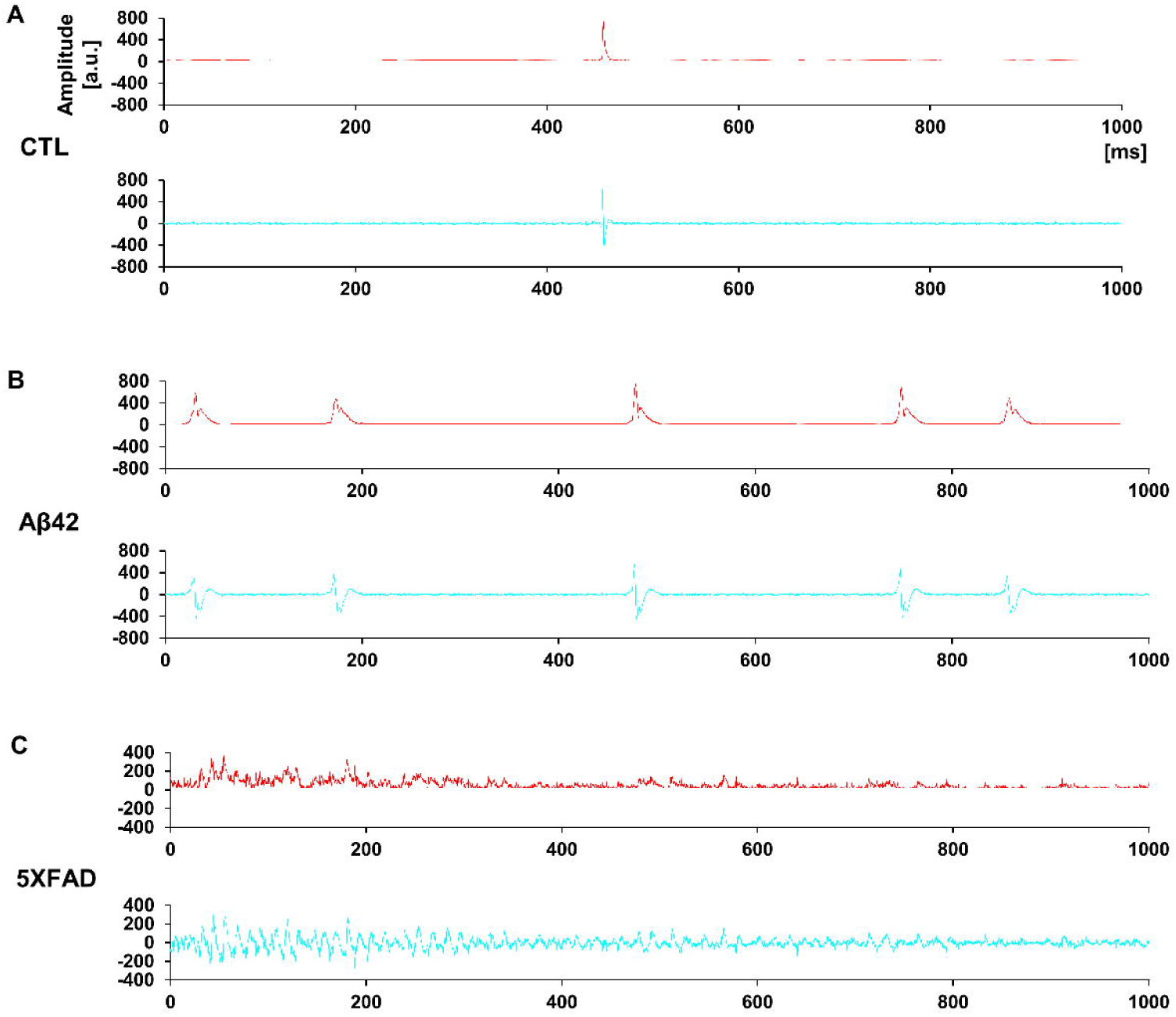
(A) CSD waveform amplitude over time separated in sources (red) and sinks (blue) in control OHSC. (B) CSD waveform amplitude over time separated in sources (red) and sinks (blue) in Aβ42 OHSC. (C) CSD waveform amplitude over time separated in sources (red) and sinks (blue) in 5XFAD NLF exosome-treated OHSC.

**MOVIE 1,2,3** CSD amplitude changes over time in 3D graph and geographical location of control (MOV.1), Aβ42 (MOV.2), and 5XFAD NLF exosome treated OHSC (MOV.3). Red for sources and blue for sinks.

## Acknowledgement

This research was supported by a grant of the Korea Health Technology R&D Project through the Korea Health Industry Development Institute (KHIDI), funded by the Ministry of Health & Welfare, Republic of Korea (grant number: HI17C1711). In addition, it was supported by the Bio & Medical Technology Development Program of the National Research Foundation (NRF) funded by the Ministry of Science & ICT (2018M3A9H1023323) and also supported by the National Research Foundation of Korea (NRF) grant funded by the Korea government (MSIT) (2022R1A2C1093134). We thank Dr. Hyun Suk of Kangwon National University for his help in the preparation of TEM images. We also thank GENUV Inc. for providing 5XFAD animals.

## Notes

### Competing Interest Statement

The authors have declared no competing interest.

### Summary of Updates

1. Figure 3, 4, 5 revised and the corresponding result descriptions 2. Introduction revised 3. Method revised

